# Δ9-tetrahydrocannabinol inhibits Hedgehog-dependent patterning during development

**DOI:** 10.1101/2021.01.18.427140

**Authors:** Hsiao-Fan Lo, Mingi Hong, Henrietta Szutorisz, Yasmin L. Hurd, Robert S. Krauss

**Affiliations:** Department of Cell, Developmental, and Regenerative Biology, Icahn School of Medicine at Mount Sinai, New York, NY 10029, USA; Addiction Institute and Departments of Psychiatry and Neuroscience, Icahn School of Medicine at Mount Sinai, New York, NY 10029, USA

**Author notes:** Corresponding Author: Robert S. Krauss, Department of Cell, Developmental, and Regenerative Biology, Icahn School of Medicine at Mount Sinai, New York, NY 10029.

**Keywords:** Δ9-tetrahydrocannabinol, Hedgehog, holoprosencephaly, birth defect, cannabis

## Abstract

Many birth defects are thought to arise from a multifactorial etiology; i.e., interaction between genetic and environmental risk factors. The Hedgehog (HH) signaling pathway regulates myriad developmental processes, and pathway inhibition is associated with birth defects, including holoprosencephaly (HPE). Cannabinoids are HH pathway inhibitors, but little is known of their effects on HH-dependent processes in mammalian embryos, and their mechanism of action is unclear. We report here that the psychoactive cannabinoid Δ9-tetrahydrocannabinol (THC) induces two hallmark HH loss-of-function phenotypes (HPE and ventral neural tube patterning defects) in *Cdon* mutant mice, which have a subthreshold deficit in HH signaling. THC therefore acts as a “conditional teratogen”, dependent on a complementing but insufficient genetic insult. In vitro findings indicate that THC is a direct, albeit relatively weak, inhibitor of the essential HH pathway component, Smoothened. In contrast, the canonical THC receptor, cannabinoid receptor-type 1, is not required for THC to inhibit HH signaling. Cannabis consumption during pregnancy may contribute to the combination of risk factors underlying specific developmental disorders. These findings therefore have significant public health relevance.

## Introduction

Congenital malformations affect approximately 8 million newborns each year and are a leading cause of death for infants and children of all ages (Christianson *et al*, 2006; Krauss & Hong, 2016; Wallingford, 2019). In some cases, mutations in single genes or exposure to individual teratogens is sufficient to cause a developmental disorder in most or all those affected (Amberger *et al*, 2019; Gilbert-Barness, 2010; Webber *et al*, 2015). For many of the most common structural birth defects, however, a single causative factor cannot be identified, and the underlying etiology for such disorders is poorly understood. In these cases, it is likely that genetic and environmental risk factors interact to elevate the chance of a defect occurring in specific developmental processes (Beames & Lipinski, 2020; Fraser, 1980; Krauss & Hong, 2016; Lovely *et al*, 2017). Genome sequencing has led to identification of numerous birth defect-associated variants, many of which appear to predispose to a given anomaly and presumably act with additional factors (Webber *et al*., 2015). Identification of subthreshold environmental risk factors by epidemiology is more difficult.

Cannabis is the illicit drug most commonly used during pregnancy and, with expanded legalization and decreased perception of risk, use is increasing (Volkow *et al*, 2017; Young-Wolff *et al*, 2017). Meta-analysis of studies through 2014 concluded that maternal cannabis use is associated with low birth weight and increased likelihood of NICU placement (Gunn *et al*, 2016). Recently, in Colorado, a correlation was reported between: 1) increased cannabis usage during pregnancy; 2) increased fetal phytocannabinoid exposure levels; and 3) an increase in major structural developmental defects (Reece & Hulse, 2019). Use of other drugs and tobacco remained static or fell in Colorado during the reporting period. A similar pattern was observed with rising incidence of atrial septal defects in multiple US states and Australia (Reece & Hulse, 2020). These correlations suggest that cannabinoids might be teratogenic, but they do not demonstrate causality.

A common birth defect that serves as a model for gene-environment interactions and multifactorial etiology is holoprosencephaly (HPE) (Beames & Lipinski, 2020; Hong & Krauss, 2018; Roessler *et al*, 2018). HPE is caused by failure to define the midline of the forebrain and/or midface. HPE comprises a phenotypic continuum ranging from complete failure to partition the forebrain into hemispheres with accompanying cyclopia, through to mild midfacial midline deficiency (Cohen, 2006; Muenke & Beachy, 2001). The Hedgehog (HH) signaling pathway is a key regulator of many developmental processes, including patterning of the forebrain and facial midline, limbs and digits, and ventral neural tube (VNT) (Petryk *et al*, 2015; Sagner & Briscoe, 2019; Tickle & Towers, 2017). HPE is associated with heterozygous mutations in the HH pathway (Dubourg *et al*, 2018; Roessler *et al*., 2018). Clinical presentation of HPE is highly variable, and many mutation carriers are unaffected, even in pedigrees. These observations have led to a multifactorial, “mutation plus modifier” model, in which heterozygous mutations may be insufficient for severe phenotypes and their penetrance and expressivity are graded by additional genetic variants and/or environmental exposures (Dubourg *et al*., 2018; Hong & Krauss, 2018; Roessler *et al*, 2012).

HH ligands activate a conserved signaling pathway (Kong *et al*, 2019; Lee *et al*, 2016; Petrov *et al*, 2017). In the absence of HH, the primary receptor Patched1 (PTCH1) acts to inhibit the activity of a second membrane protein, Smoothened (SMO). Binding of HH to PTCH1 relieves inhibition of SMO. SMO then signals to activate pathway target genes via GLI transcription factors. SMO is a class F, G protein-coupled receptor (GPCR). PTCH1 appears to function as a transporter to restrict accessibility of SMO to its activating ligands, namely cholesterol and oxysterols. HH ligands block PTCH1 function, allowing SMO access to cholesterol and oxysterols, thus activating SMO signaling (Qi & Li, 2020; Radhakrishnan *et al*, 2020). These events occur in the primary cilium, an organelle in which HH pathway components traffic and concentrate for signaling (Bangs & Anderson, 2017; Gigante & Caspary, 2020). Consistent with the notion that SMO is itself a ligand-regulated receptor, many small molecules that act as SMO agonists or antagonists have been identified (Sharpe *et al*, 2015). While PTCH1 function is sufficient for SMO inhibition, HH signal reception also requires at least one of three coreceptors (CDON, BOC, GAS1). CDON, BOC, and GAS1 have overlapping roles and are collectively required for HH signaling (Allen *et al*, 2011; Izzi *et al*, 2011; Wierbowski *et al*, 2020; Zhang *et al*, 2011). Mice with targeted mutations in any one of these coreceptors have a selective and partial loss of HH pathway function.

In a search for endogenous lipids that act as SMO antagonists, Eaton and colleagues identified endocannabinoids as inhibitors of HH signaling (Khaliullina *et al*, 2015). Endocannabinoids were effective as HH pathway inhibitors in both developing *Drosophila* wing disks and cultured mammalian cells. Endocannabinoids are fatty acids/alcohols linked to polar head groups that signal through the GPCRs, cannabinoid receptor-type 1 (CB1R) and −type 2 (CB2R) (Lu & Mackie, 2020; Maccarrone *et al*, 2015). Phytocannabinoids are bioactive ingredients of cannabis and include Δ^9^-tetrahydrocannabinol (THC) and cannabidiol (CBD), the former being the major psychoactive component. These compounds exert their effects via CB1R, CB2R, and/or other receptors, with CB1R responsible for mediating the major neurobehavioral effects of THC (Lu & Mackie, 2020; Schurman *et al*, 2020). Importantly, THC and CBD inhibited HH signaling similarly to endocannabinoids, whereas structurally unrelated CB1R and CB2R agonists/antagonists did not (Khaliullina *et al*., 2015). Cannabinoids appeared to inhibit HH signaling at the level of SMO, although this mechanism was not uniform among those analyzed and has been questioned by others (Sever *et al*, 2016).

These findings raise the possibility that in utero exposure to phytocannabinoids might be teratogenic, via an ability to inhibit HH signaling at critical points during development. We report here that THC is teratogenic to genetically-sensitized mice harboring a subthreshold deficit in HH pathway signaling strength. THC dose-dependently induced HH loss-of-function phenotypes in these mice, including HPE, but not in wild type mice. THC therefore acted as a “conditional teratogen”, dependent on a complementing but insufficient genetic insult. Furthermore, in *in vitro* assays, THC displayed properties similar to a *bone fide* SMO inhibitor, SANT-1. Together, these results raise the possibility that human cannabis consumption during early pregnancy may expose embryos to a HH inhibitor and environmental risk factor for birth defects.

## Results

### THC-exposed *Cdon*^-/-^ embryos display HPE with craniofacial midline defects

*Cdon*^-/-^ mice on a 129S6 background have a subthreshold defect in HH signaling and are sensitive to induction of HPE by both genetic and environmental modifiers (Hong & Krauss, 2018). They therefore model human HPE and are an ideal model to assess THC teratogenicity. 129S6 *Cdon^+/-^* mice were intercrossed and pregnant females received a single dose of THC at 5, 10, or 15 mg/kg, administered IP at E7.5. The E7.5 time point was used because it was most effective for induction of HPE in wild type mice by the potent SMO inhibitor Vismodegib (Heyne *et al*, 2015). We measured blood concentrations of THC and metabolites in pregnant female mice, and the peak levels achieved at the highest THC dose used were similar to those achieved by humans after inhalation of 34mg cannabis (180-200 ng/ml) (Grotenhermen, 2003) (Supplemental Fig. 1).

Embryos were harvested at E14 and scored for HPE phenotypes in whole mount and thin sectioned samples. Vehicle- and THC-treated *Cdon*^+/+^ and *Cdon*^+/-^ embryos did not have HPE, nor did vehicle-treated *Cdon*^-/-^ embryos. In contrast, THC-exposed *Cdon*^-/-^ embryos displayed mid-facial HPE phenotypes, including fused upper lip, in a dose-responsive manner (Figs. 1A, 1B, Supplemental Table 1). H&E-stained sections revealed that THC-treated *Cdon*^-/-^ embryos had fully partitioned forebrains, but displayed substantial diminution of midfacial structures, including close-set and rudimentary vomeronasal organs and pronounced loss of midfacial midline structure (Fig. 1B). These results classify THC-induced HPE in *Cdon^-/-^* mice as of the relatively mild, lobar category (Krauss, 2007). Whole mount in situ hybridization at E10 revealed reduction in expression of a direct SHH target gene, *Gli1* in the rostroventral midline of THC-treated *Cdon^-/-^* embryos (Fig. 1C, 1D). Expression of a second target gene, *Nkx2.1*, was also reduced THC-treated *Cdon^-/-^* embryos although variability between samples resulted in P = 0.07 (Fig. 1C, 1D). This variability may be related to the partial penetrance of the HPE phenotype. Therefore, *Cdon* mutation and THC synergized to induce HPE in mice.

**Fig. 1.**
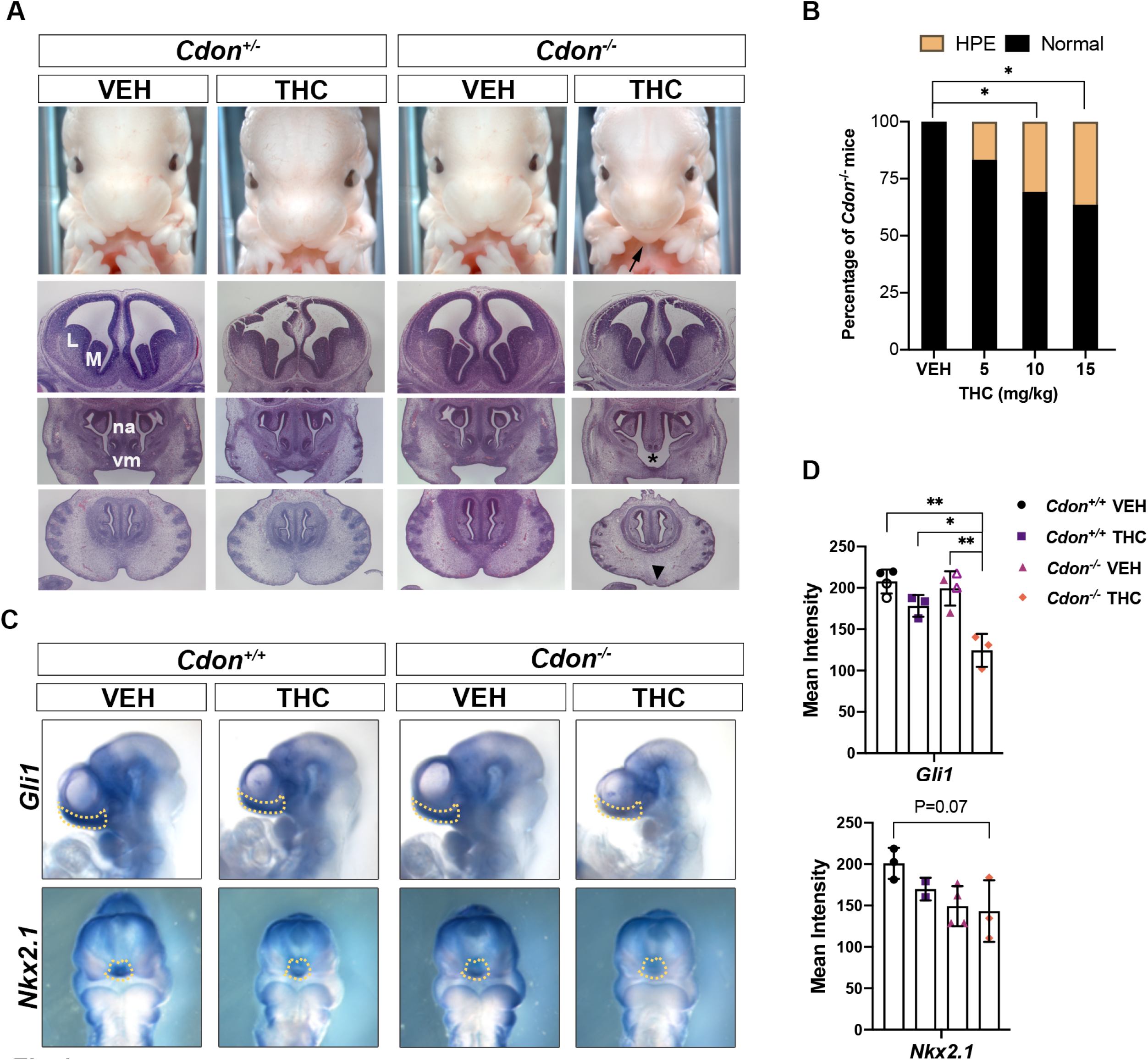
THC induces HPE in *Cdon*^-/-^ mutant mice. A. Frontal views of forebrains and faces of embryos with indicated genotypes and treatments. HPE phenotypes in THC-treated *Cdon*^-/-^ mice include fusion of the upper lip (arrow), close-set and rudimentary vomeronasal organs (vm, asterisk), and loss of midfacial midline structure (arrowhead). Top row, whole mount E14 embryos. Next three rows, H&E-stained sections of E14 embryos. VEH, vehicle-treated embryos; M, medial ganglionic eminence; L, lateral ganglionic eminence; na, nasal septum. B. THC induces HPE in *Cdon*^-/-^ mice in a dose-dependent manner. See Expanded View Table 1 for complete quantification of mice. *, p<0.05 with two-tailed Fischer’s exact test. C. Whole mount in situ hybridization of E10 embryos exposed in utero to THC (15mg/kg) or VEH. Expression of two direct SHH target genes (*Gli1* and *Nkx2.1*) is reduced in the rostroventral midline (arrows). D. Quantification of in situ hybridization. The mean intensity of the in situ hybridization signal was determined within the dashed lines of individual embryos (as shown in panel C). For the VEH conditions for *Gli1*, closed symbols are with 0.3% Tween-80 in saline (the vehicle for THC), the open symbols are with saline alone, to demonstrate the Tween-80 had no effect on its own. *, ** p<0.05, <0.01 with Student’s t-test.

### THC induces VNT patterning defects in *Cdorf*^-/-^ mice

We next sought to assess the effects of in utero exposure to THC on a second HH-dependent patterning process. Sonic HH (SHH) produced by the notochord and floor plate (FP) forms a ventral-to-dorsal gradient of pathway activity in the developing neural tube. In response to distinct levels of SHH pathway activity, expression of specific transcription factors is induced in specific progenitor zones of the VNT. These proteins include: FOXA2 (in the FP), NKX2.2 (in p3 progenitors), and OLIG2 (in pMN motor neuron progenitors). Pregnant dams were treated with a single dose of THC (15 mg/kg) at E8.0 and embryos were analyzed at E9.5, by immunofluorescence (IF) analysis of forelimb-level sections. The E8.0 time-point was used because it was effective for inhibition of SHH-dependent VNT pattering in wild type embryos by the SMO inhibitor cyclopamine (Ribes *et al*, 2010). THC reduced the number of FOXA2^+^ FP cells by >50% and of NKX2.2^+^ p3 progenitors by >35% in *Cdon^-/-^* embryos (Fig. 2A, 2B). The more dorsally positioned OLIG2^+^ pMN progenitors, which require a lower level of HH signal to be induced, were not affected (Fig. 2A, 2B). These results are very similar to those obtained by removal of one copy of *Shh* from *Cdon^-/-^* mice (Tenzen *et al*, 2006). Taken together, our findings demonstrate that THC is teratogenic in genetically sensitized mice, producing two well-established, SHH loss-of-function phenotypes: HPE and defective VNT patterning.

**Fig. 2.**
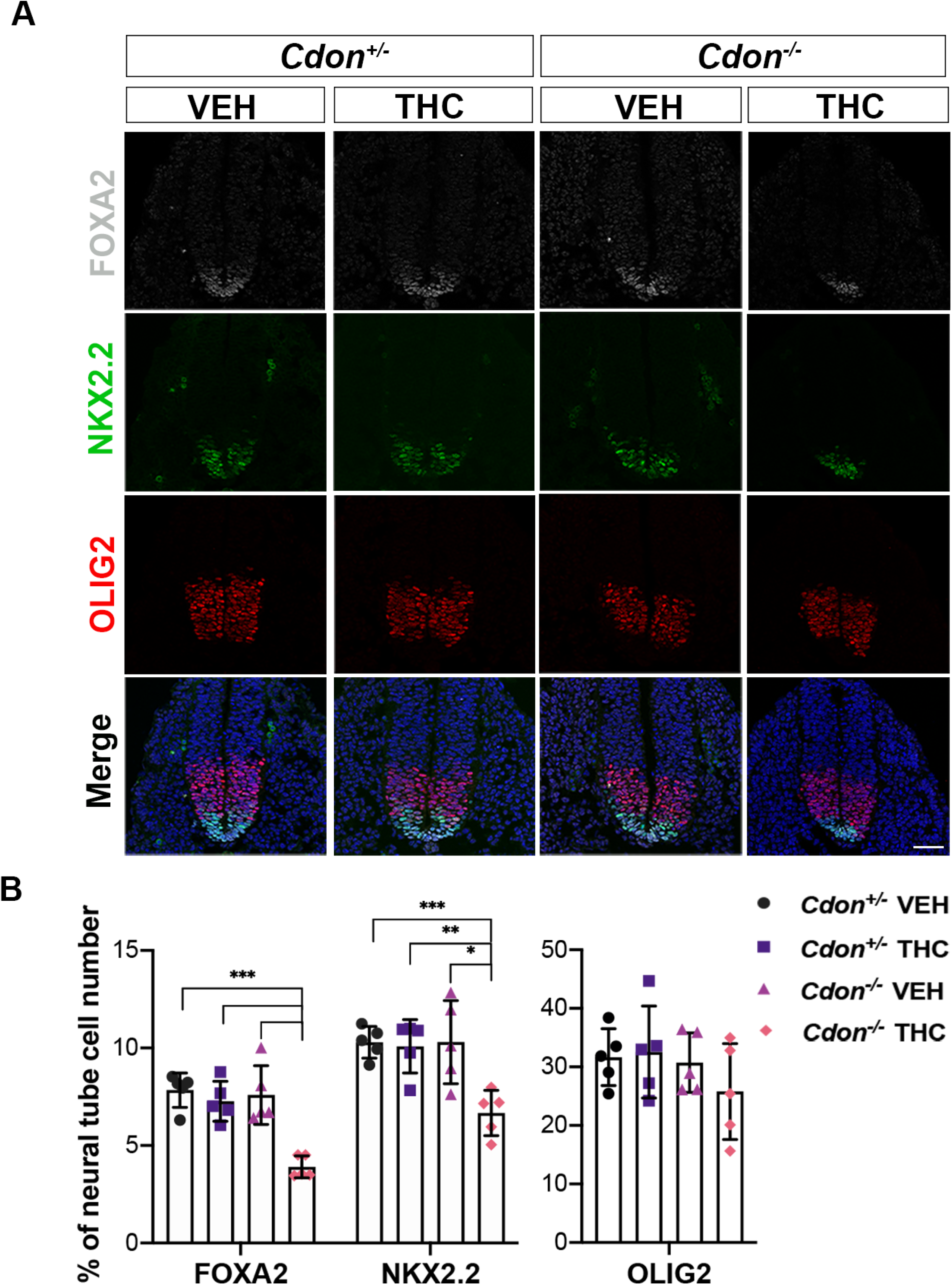
THC induces VNT patterning defects. A. SHH-dependent VNT cell types identified by immunofluorescence in sections of embryos with indicated genotypes treated in utero with THC (15mg/kg) or VEH. B. Numbers of FOXA2^+^, NKX2.2^+^ and OLIG2^+^ cells relative to whole neural tube cells were quantified. All nuclei are visible by DAPI stain (blue) in the merged images. Values are means from 3-5 sections from 5 individual mice *, **, *** p<0.05, <0.01, <0.001 with Student’s t-test. Scale bar, 50μm.

### THC is likely a direct inhibitor of SMO

Khaliullina et al. reported that THC inhibited a HH-dependent reporter construct in NIH3T3 cells, a well-established cell culture system for studying HH signaling and the mechanisms of pathway inhibitors (Khaliullina *et al*., 2015; Taipale *et al*, 2000). We confirmed this observation. Treatment of NIH3T3 cells with recombinant SHH induced an ~10-fold increase in expression of the direct, endogenous target gene, *Gli1*, as measured by qRT-PCR (Fig. 3A). THC dose-dependently inhibited *Gli1* induction in response to SHH, with an IC_50_ of ~1 μM. Similar results were obtained when these cells were stimulated with the direct SMO agonist, SAG (Supplemental Fig. 2A), and with SHH treatment of a second cell system, freshly prepared mouse embryo fibroblasts (MEFs) (Supplemental Fig. 2B).

**Fig. 3.**
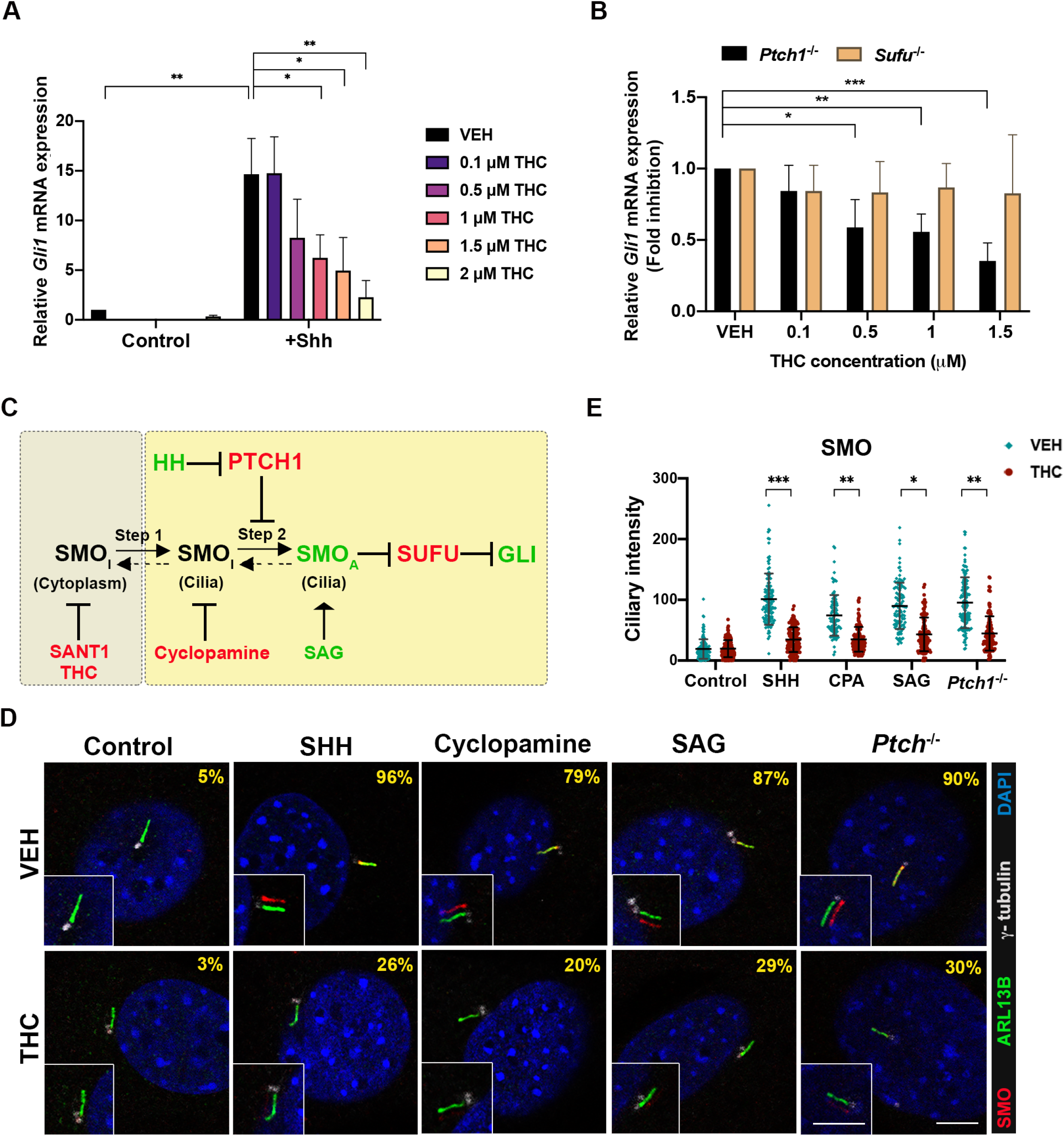
THC inhibits HH signaling at the level of SMO. A. NIH3T3 cells were treated with 5 ng/ml SHH and the indicated concentrations of THC for 24 hr. Relative endogenous *Gli1* mRNA expression was analyzed by qRT-PCR and normalized to *Gapdh* levels. Values are means ± SD, N=3. B. THC inhibits endogenous *Gli1* expression in *Ptch1^-/-^*, but not *Sufu^-/-^* MEFs. Relative *Gli1* mRNA expression was determined by qRT-PCR and normalized to *Gapdh* levels. Values are means ± SD, N=3. C. SMO activation is a two-step process: ciliary transport of inactive SMO (SMO_I_) (step 1) is followed by SMO activation (SMO_A_) within cilia (step 2). Pathway agonists (green) and antagonists (red) can control SMO transport entry or exit steps. D. THC blocks endogenous SMO localization in primary cilia in NIH3T3 cells in response to multiple stimuli. Primary cilia are marked by ARL13B, γ-tubulin marks centrioles. The percentage of cells with cilium-localized SMO is displayed in the upper right. See Expanded View Table 2 for complete quantification. Inset: The SMO and ARL13B signal overlay is shifted for easy visualization. SHH was used at 5ng/ml, cyclopamine at 5μM, SAG at 100nM, and THC at 1.5μM. Scale bar, 5μm. E. Quantification of SMO fluorescence intensity in primary cilia. Each point represents an individual cell collated from at least 3 independent experiments. *, **, *** p<0.05, <0.01, <0.001 with Student’s t-test.

It was reported that cannabinoids inhibited HH signaling at the level of SMO, although differences were seen between the cannabinoids examined, and THC was not tested (Khaliullina *et al*., 2015). PTCH1 and SUFU are negative regulators of HH signaling. MEFs that are null for either gene display constitutive pathway activity, with PTCH1 functioning upstream of SMO, and SUFU acting downstream of SMO (Kong *et al*., 2019; Lee *et al*., 2016; Petrov *et al*., 2017). To identify a position within the HH pathway at which THC acts, we first tested its ability to inhibit constitutive expression of *Gli1* in *Ptch1^-/-^* and *Sufu*^-/-^ MEFs. THC attenuated *Gli1* expression in *Ptch*^-/-^ MEFs, but not in *Sufu*^-/-^ MEFs, in a dose-dependent manner (Fig. 3B). Therefore, THC modulates SHH signaling downstream of PTCH1 and upstream of SUFU. While these data place THC’s likely point of action at the level of SMO, this could occur via direct SMO inhibition or indirectly, e.g., via perturbation of primary cilia, the cellular site of SMO signaling.

SMO activation is a two-step process. Ciliary transport of SMO (step 1) is followed by SMO activation within cilia (step 2) (Fig. 3C) (Rohatgi *et al*, 2009; Wilson *et al*, 2009). A low level of SMO traffics through primary cilia constitutively. Some direct SMO inhibitors, like SANT-1, induce a SMO conformation that inhibits trafficking and prevents ciliary accumulation in response to activators, such as SHH and SAG. Another class of SMO inhibitor, exemplified by cyclopamine, induces a SMO conformation that does not inhibit trafficking, but which resists activation within cilia. Therefore, cyclopamine-type inhibitors trap SMO in primary cilia even in the absence of SHH or SAG, yet block activation of SMO within cilia, even in the presence of these pathway activators (Rohatgi *et al*., 2009; Wilson *et al*., 2009). We assessed THC’s action on ciliary translocation of endogenous SMO in NIH3T3 cells treated with various HH pathway-regulating compounds (Fig. 3D, 3E, Supplemental Table 2). The percentage of cells with ciliary SMO and fluorescent intensity of ciliary protein expression were determined. THC alone did not trigger SMO translocation to cilia, but it blocked SMO accumulation in cilia in response to SHH, cyclopamine, or SAG. THC also inhibited constitutive ciliary localization of SMO in *Ptch*^-/-^ MEFs. THC did not affect cilium length or ciliary levels of the primary cilia marker, ARL13B (Supplemental Fig. 3A, 3B). Therefore, THC acted similarly to SANT-1, preventing SMO translocation into primary cilia.

The phytocannabinoid CBD is structurally related to THC and was also reported to inhibit HH signaling (Khaliullina *et al*., 2015). In contrast to our results with THC, Khaliullina et al. found that CBD did not to prevent localization of exogenously expressed SMO to primary cilia in response to SAG (Khaliullina *et al*., 2015). Because these results and our own findings with THC were dichotomous, we tested CBD as well. CBD inhibited SHH-induced *Gli1* expression in NIH3T3 cells with an IC_50_ of ~1.2μM, a value similar to that of THC (Fig. 4A). Like THC, CBD alone did not promote translocation of SMO to primary cilia (Fig. 4B, 4C). We found that CBD did reduce endogenous SMO ciliary translocation in response to SHH. However, despite having similar IC50 values, CBD was significantly less effective than THC in reducing: 1) the percentage of cells with SMO^+^ cilia; and 2) the steady-state amount of SMO within SMO^+^ cilia, in SHH-treated cells (Fig. 4B, 4C, Supplemental Table 2). CBD did not affect cilium length or ciliary levels of the primary cilia marker, ARL13B (Fig. 4C). CBD may therefore inhibit HH signaling by a mechanism somewhat distinct from either SANT-1-type or cyclopamine-type SMO inhibitors.

**Fig. 4.**
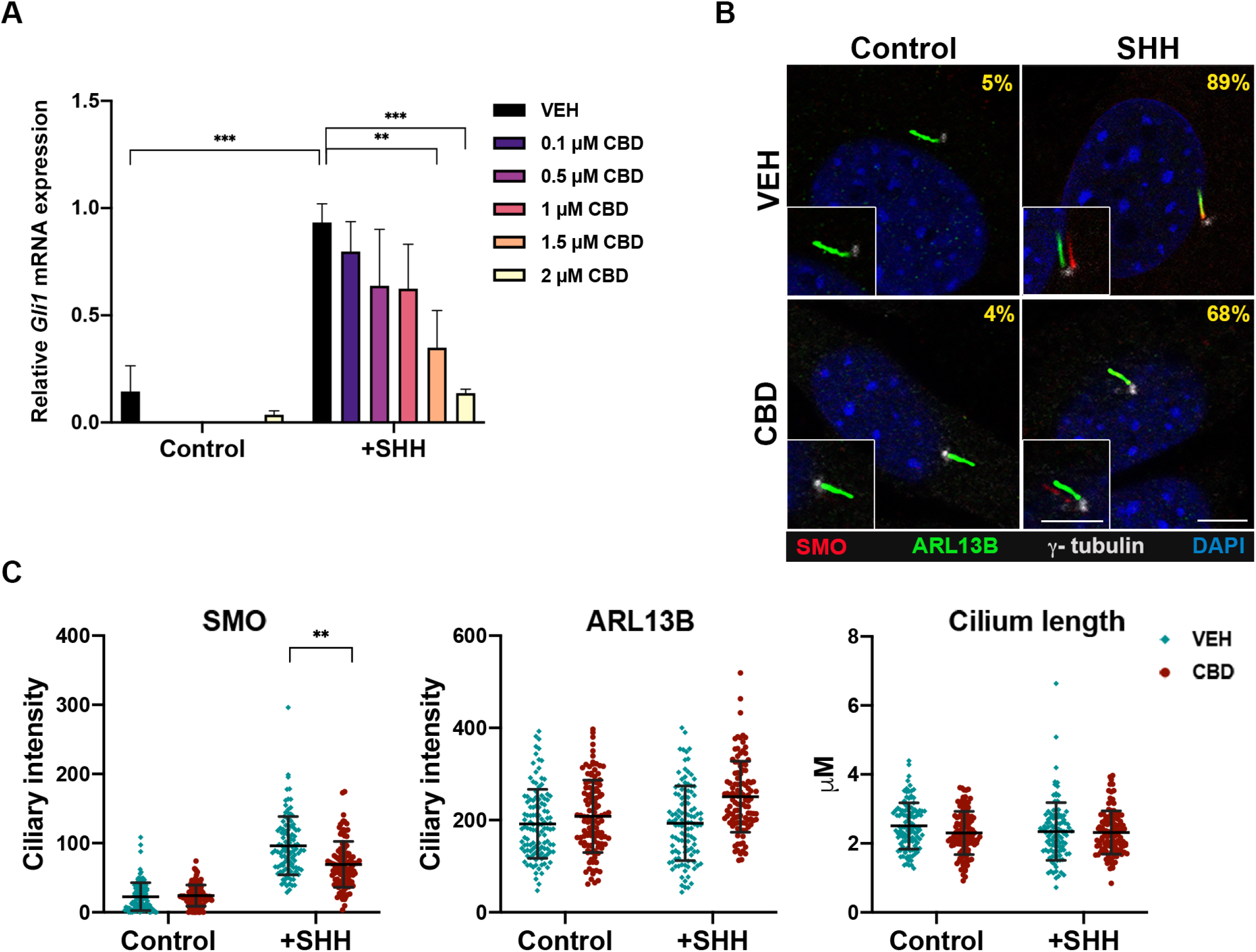
CBD inhibits SHH pathway activation and reduces SMO localization in primary cilia. A. CBD dose-dependently inhibited Gli1 expression with SHH stimulation in NIH3T3 cells. B. The percentage (upper right of each panel) of SMO translocate into cilia in SHH-stimulated cells was modestly decreased by CBD. See Expanded View Table 2 for complete quantification. Inset: The SMO and ARL13B signal overlay is shifted for easy visualization. SHH was used at 5ng/ml and CBD at 1.5μM. Scale bar, 5μm. C. Quantification of SMO and ARL13B fluorescent intensity in primary cilia. Cilia length were also quantified. Each point represents an individual cell collated from 3 different experiments.

To identify whether THC may be a direct inhibitor of SMO, we measured its ability to compete with a fluorescent derivative of cyclopamine (Bodipy-CPA) for binding to SMO (Chen *et al*, 2002). We transfected HEK293 cells with an expression vector encoding Myc epitope-tagged SMO or with an empty vector as a control. SMO-Myc – expressing cells, but not control cells, displayed strong Bodipy-CPA binding (Fig. 5A, 4B). SANT-1 and KAAD-cyclopamine (a potent analog of cyclopamine) each competed with Bodipy-CPA for SMO binding (Fig. 5A, 5B). THC at 10μM, but not 1μM, also inhibited Bodipy-CPA binding (Fig. 5A, 5B). In contrast, 10μM CBD did not significantly diminish Bodipy-CPA binding to SMO-expressing cells (Fig. 5A, 5B). While 10μM is a high concentration of THC, cholesterol (an endogenous SMO ligand) competed with Bodipy-CPA in a similar concentration range (Huang *et al*, 2016). Furthermore, the fact that THC’s close structural congener CBD was not effective at this dose argues that THC’s ability to compete with Bodipy-CPA was specific.

**Fig. 5.**
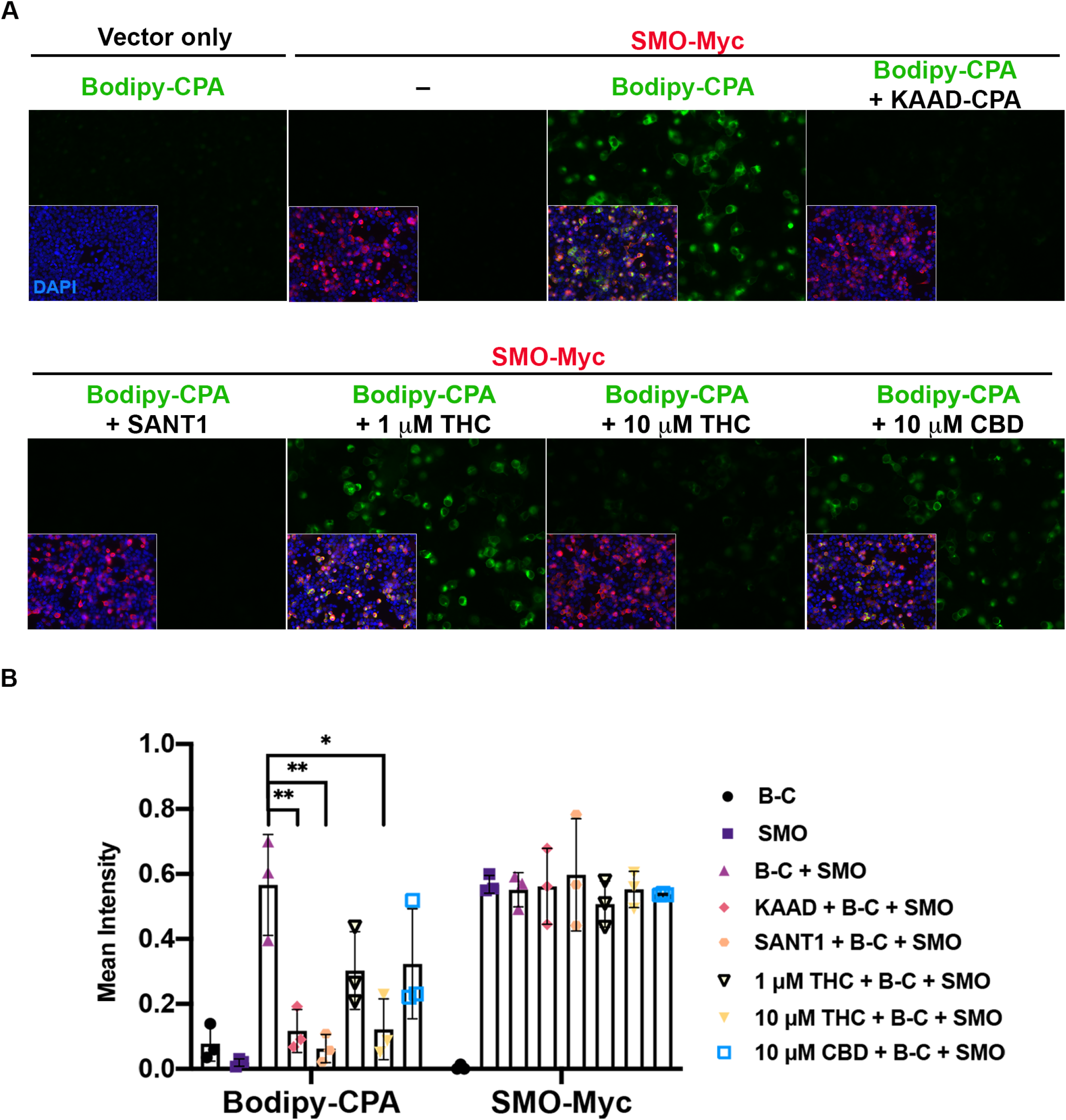
THC, but not CBD, competes with Bodipy-cyclopamine for binding to SMO. A. HEK293T cells were transfected with SMO-Myc (red, insets) or empty vector. Cells were then incubated with Bodipy-cyclopamine (Bodipy-CPA; green), in the presence or absence of different competitors as indicated. Bodipy-CPA was used at 5nM, KAAD-CPA at 200nM, SANT 1 at 1μM, and THC and CBD as indicated. B. Quantification of Bodipy-CPA (B-C) and SMO-Myc fluorescence intensity. *, ** p<0.05, <0.001, with Student’s t-test. Values represent means ± SD, from three independent experiments.

### CB1R is not required for THC-mediated inhibition of HH signaling

THC’s psychoactive effects are exerted via CB1R, encoded by the gene *Cnr1* (Lu & Mackie, 2020). It has been proposed that cannabinoids inhibit HH signaling via CB1R-SMO heterodimers (Fish *et al*, 2019). We sought to explore this possibility further. First, if CB1R is required for cannabinoids to inhibit SMO, CB1R must be expressed in cells that display a cannabinoid-sensitive HH response. NIH3T3 cells and MEFs are HH responsive, and THC inhibits this response (Fig. 3A, Fig. S2A, S2B). However, we did not detect *Cnr1* mRNA and or CB1R protein in either cell type, whereas they were easily detected in mouse whole brain lysate (Fig. 6A, 6B).

**Fig. 6.**
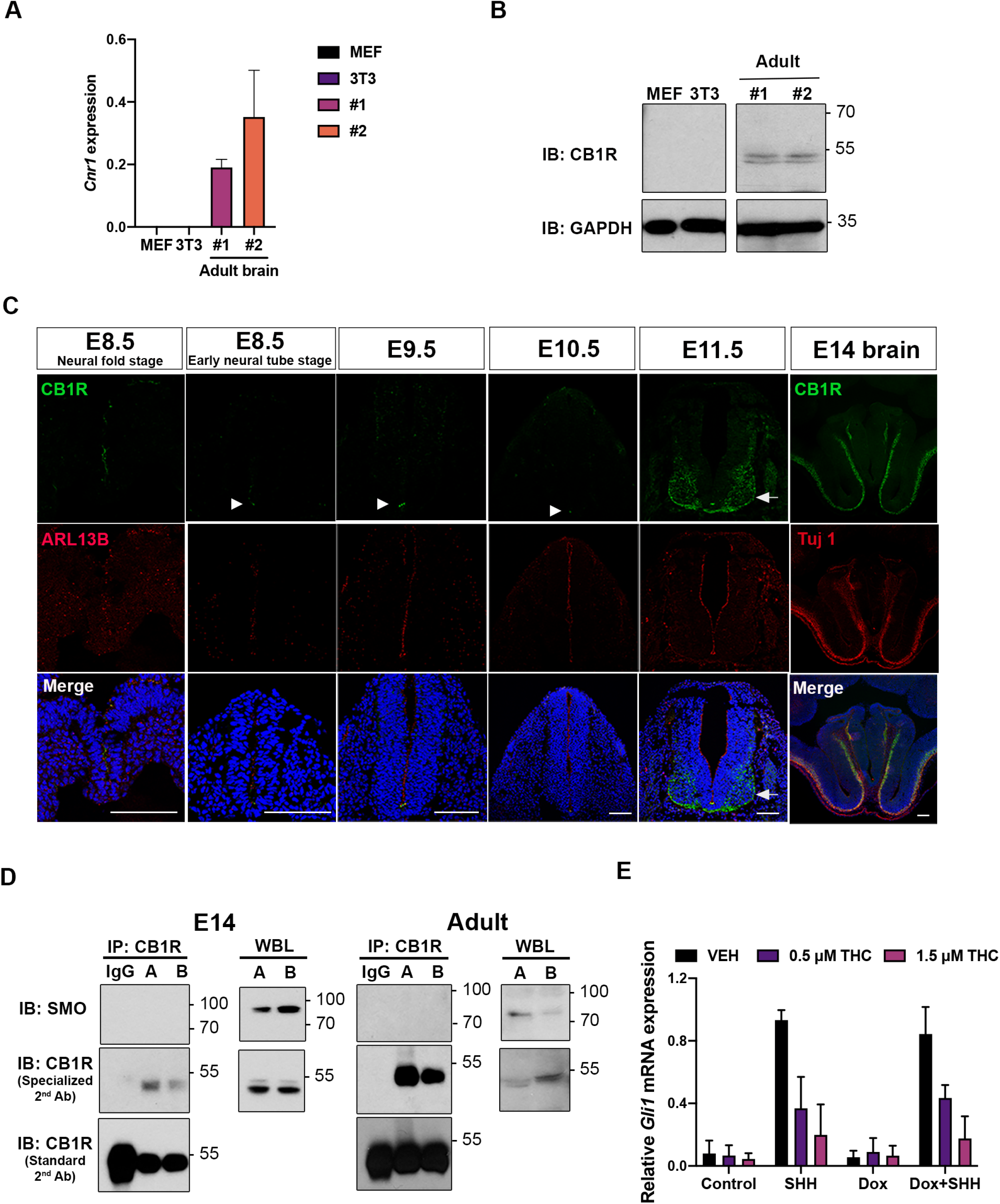
CB1R is not required for THC-mediated inhibition of HH signaling. A and B. Expression of *Cnr1* mRNA in MEFs, NIH3T3 cells, and adult mouse brain (2 different samples) by qRT-PCR (A) and immunoblotting (B). Note that HH-responsive MEFs and NIH3T3 cells do not express *Cnr1* /CB1R. Number of PCR cycles: MEFs=30; 3T3=35; brain=21. C. Forelimb level sections of mouse neural tubes at the indicated stages were stained with antibodies to CB1R and ARL13B and with DAPI (blue). Little to no CB1R is detected in the neural tube at E8.5, E9.5, or E10.5, except in a small number of cells in the ventricular zone of the floor plate (arrowheads). CB1R expression in the non-HH-responsive mantle zone at E11.5 is indicated by the arrow. ARL13B marks ciliated cells in the HH-responsive ventricular zone. CB1R staining of an E14 mouse brain section is used as a control, along with the neural differentiation marker, TuJ1 (red). Scale bars, 100μm. D. SMO does not co-immunoprecipitate (IP) with CB1R in lysates of E14 mouse brain or adult mouse brain (two independent samples, designated A and B for the former and 1 and 2 for the latter). Whole brain lysates (WBL) were directly immunoblotted (IB) as a control. Light chain specific (GeneTex or Jackson Lab) secondary antibody (“specialized 2^nd^ Ab) was used to avoid detection of IgG heavy chain, which is of similar MW to CB1R (and seen in the blots using a “Standard 2nd Ab”. The IgG lane is an IP with control, non-immune IgG in place of anti-CB1R antibody. E. Exogenous expression of CB1R in NIH3T3 cells does not alter THC dose-dependent inhibition of SHH signaling, measured by induction of endogenous *Gli1* expression. Dox, cells pretreated with doxycycline to induce CB1R expression (see Fig. EV4A). Values are means ± SD, N=3.

We next performed IF analysis for CB1R on VNT sections from embryos collected at E8.5 (both neural fold and early NT stages), E9.5, E10.5, and E11.5. CB1R was detected in some cells of the ventricular zone at the neural fold stage, consistent with previous reports (Gilbert *et al*, 2015; Psychoyos *et al*, 2012). However, little to no specific CB1R IF signal was observed in the developing VNT at E8.5 early neural tube stage, E9.5, or E10.5, with the exception of a few cells at the ventricular edge of the floor plate that reproducibly gave strong signal (Fig. 6C). Robust CB1R expression was observed at E11.5, but this was localized to the HH-non-responsive mantle domain, which contains more differentiated neurons (Fig. 6C). These results demonstrate that CB1R levels are very low in VNT cells at developmental stages when they are undergoing THC-sensitive, SHH-dependent neuronal patterning. These findings are consistent with mRNA in situ hybridization data, showing a lack of *Cnr1* expression at similar stages in the chick VNT (Watson *et al*, 2008), patterning of which closely resembles the mouse VNT (Dessaud *et al*, 2008). In general, *Cnr1* expression follows neuronal differentiation (Watson *et al*., 2008), consistent with our IF results in the VNT. Our IF with CB1R antibodies was further validated by staining sections of E14 cerebral cortex (Fig. 6C), which reproduced similar, previously reported localization findings (Mulder *et al*, 2008).

We next attempted to co-immunoprecipitate CB1R and SMO. Extracts of E14 and adult mouse brains were immunoprecipitated with antibodies to CB1R and then blotted for CB1R and SMO. CB1R and SMO were detected in whole lysates by Western blot. However, while CB1R was efficiently immunoprecipitated, SMO was not detectable in these precipitates (Fig. 6D).

Finally, to test for effects of CB1R in THC-mediated inhibition of HH signaling, we established NIH3T3 cells that expressed CB1R in a doxycycline-inducible manner. We reasoned that if CB1R levels were rate-limiting for THC’s ability to inhibit HH signaling, exogenous expression of CB1R in NIH3T3 cells would sensitize them to THC-mediated attenuation of SHH-induced expression of *Gli1*. Treatment of these cells with 0.05μg/ml doxycycline induced CB1R to a level similar to that seen in Neuro-2a cells, which display CB1R-dependent signaling in response to cannabinoids (Graham *et al*, 2006; He *et al*, 2005) (Supplemental Fig. 4A). CB1R expression did not alter the ability of THC to dose-dependently inhibit *Gli1* mRNA induction in response to SHH stimulation (Fig. 6E). Furthermore, CB1R expressed in NIH3T3 cells was not present in primary cilia, in control, THC-, SHH-, or THC+SHH-treated cells (Supplemental Fig. 4B). Taken together, we conclude that CB1R is not required for THC-mediated inhibition of HH signaling.

## Discussion

We report here that THC is teratogenic in genetically sensitized mice (129S6 *Cdon^-/-^* mice). A single in utero dose of THC resulted in two hallmark HH loss-of-function phenotypes, HPE and VNT patterning defects. In vitro experiments argue that THC exerted these effects as a direct, albeit relatively weak, inhibitor of the essential HH pathway signal transducer, SMO. As THC’s effects were dependent on a complementary, but insufficient, genetic deficiency, we categorize it as a “conditional teratogen”. Cannabis consumption during pregnancy may therefore result in partial inhibition of a major morphogenetic pathway in embryos, thereby contributing to the complex combination of genetic and environmental risk factors that are thought to underlie many common developmental disorders.

### THC inhibits HH signaling in vitro and in vivo

Eaton and colleagues first reported that endocannabinoids and phytocannabinoids inhibited HH signaling, but did not investigate cannabinoid effects in a vertebrate model *in vivo*, and did not focus on THC (Khaliullina *et al*., 2015). Our results are consistent with the conclusion that THC is a direct inhibitor of SMO. These findings include: 1) THC’s inhibitory activity mapped downstream of PTCH1 and upstream of SUFU; 2) THC prevented translocation of SMO to primary cilia upon pathway activation, without obviously altering cilia themselves; and 3) THC competed with a cyclopamine derivative for binding to SMO expressed on cells. THC therefore acted similarly to the well-studied SMO inhibitor SANT-1, although it is much less potent than SANT-1. THC’s ability to induce HH loss of function phenotypes in mice with a subthreshold HH signaling defect is fully consistent with its function as a relatively weak SMO inhibitor. In fact, THC treatment and *Shh* heterozygosity (which alone does not confer a phenotype) have similar enhancing effects on these mice (Tenzen *et al*., 2006).

The THC structural analog CBD inhibited HH signaling in vitro with a similar IC50 as THC, but was much weaker at affecting SMO translocation to cilia and did not effectively compete for binding to SMO. CBD is therefore likely to inhibit HH signaling by a molecular mechanism somewhat distinct from that of THC, even if SMO might be its target. THC and CBD exert their effects outside the HH pathway via different receptors as well, so despite being closely related, these compounds have important structural features that distinguish their activities (Dos Santos *et al*, 2021). THC’s psychoactive properties are the major reason people consume cannabis; these properties are not shared by CBD. However, CBD is available over the counter and viewed by many as generally beneficial, or at least innocuous, so it will be important to test CBD for conditional teratogenicity also.

While this work was in progress, Fish et al. reported studies on the effects of several phytocannabinoids and synthetic cannabinoids, including THC, in C57BL/6J mice (Fish *et al*., 2019). Consistent with the work reported here, THC induced “minor alterations of pupil shape and size” in these mice, but exacerbated effects on eye morphology induced by fetal alcohol exposure (Fish *et al*., 2019).

### CB1R is not required for THC to inhibit HH pathway signaling

It has been suggested that cannabinoids inhibit HH signaling via CB1R-SMO heterodimers (Fish *et al*., 2019). Evidence for this conclusion included co-immunoprecipitation of CB1R with SMO (Fish *et al*., 2019). However, our findings showed that CB1R is not required for THC’s ability to inhibit the HH pathway and argue against CB1R-SMO interaction. Evidence for these conclusions includes:

1. NIH3T3 cells and MEFs responded to HH signaling in a THC-inhibitable manner, yet they do not express detectable *Cnr1* mRNA or CB1R protein. Furthermore, THC inhibited VNT patterning, but VNT cells expressed little CB1R from E8.5-E10.5, a period when they are acquiring neuronal identity in response to SHH signaling. This is consistent with mRNA in situ hybridization for *Cnr1* expression at a similar stage of the developing chick VNT (Watson *et al*., 2008).
2. We attempted to co-immunoprecipitate CB1R and SMO without success. The previously reported, co-precipitating band identified as CB1R in anti-SMO immunoprecipitates migrated at ~100 kDa (Fish *et al*., 2019). The calculated MW of CB1R is 53 kDa, approximately the size of the band we detected in our experiments. A recent study used multiple CB1R antibodies against brain lysates from control and *Cnr1* null mice (Esteban *et al*, 2020). All antibodies recognized a band of ~53 kDa, that was specifically lost in the KO mice. It is likely, therefore, that the ~100 kDa band that co-immunoprecipitated with SMO was not CB1R.
3. Exogenous expression of CB1R in NIH3T3 cells did not alter dose-dependency of THC-mediated inhibition of SHH signaling. Furthermore, CB1R did not localize to primary cilia in the presence or absence of THC, plus or minus SHH ligand. It was reported that the CB1R inhibitor SR141716A rescued eye phenotypes associated with in utero exposure of C57BL/6 mice to the synthetic cannabinoid, CP55,940 or to CP55,940 plus alcohol (Fish *et al*., 2019). However, the eye phenotypes in this system are not distinctive to HH loss of function, and CB1R agonism may play a role in their genesis.
4. Although the role of endocannabinoids as inhibitors of vertebrate HH signaling is unknown, it is very unlikely that any such role would require CB1R or CB2R, as *Cnr1;Cnr2* double mutant mice are viable and do not display any reported HH gain-of-function phenotypes (Rowley *et al*, 2017; Sophocleous *et al*, 2017). This contrasts with strong gain-of-function phenotypes seen in embryos upon loss of established negative regulators of the HH pathway (e.g., *Ptch1, Sufu, Gnas, Gpr161, Ankmy2*) (Goodrich *et al*, 1997; Mukhopadhyay *et al*, 2013; Regard *et al*, 2013; Somatilaka *et al*, 2020; Svard *et al*, 2006).

Taken together, we conclude that CB1R is not required for THC to inhibit HH signaling. While our findings do not conclusively rule out formation of CB1R-SMO complexes, neither our work nor published studies offer compelling evidence that such complexes exist.

### THC as a potential risk factor in developmental disorders

The causes of common developmental disorders are often unknown. Many cases likely arise from a combination of subthreshold genetic and environmental insults, i.e., multifactorial etiology (Beames & Lipinski, 2020; Krauss & Hong, 2016). Whole genome sequencing should eventually reveal a great majority of genetic contributions to such defects; subthreshold environmental risk factors, however, are more difficult to identify. The HH signaling pathway plays a fundamental role during development and is involved in growth and morphogenesis of a wide variety of body structures, including limbs, brain, heart, and craniofacial structures (Petryk *et al*., 2015; Sagner & Briscoe, 2019; Tickle & Towers, 2017). Mutations in HH pathway components are involved in several developmental disorders, including Greig syndrome, Gorlin syndrome, Brachydactyly A1, and HPE (Nieuwenhuis & Hui, 2005). Environmental agents that perturb HH signaling during specific times during embryogenesis may therefore be risk factors for developmental disorders, perhaps working in concert with predisposing genetic variants.

Cannabis is the most frequently used illicit drug during pregnancy and in young women of childbearing age, and as legalization progresses, use will presumably increase. Furthermore, the THC concentration of both medicinal and recreational cannabis is currently very high (~20% THC) (Cash *et al*, 2020). Identification of a potential role of cannabis consumption in the etiology of structural birth defects is significant, as it would represent an avoidable risk factor. Cannabinoids are HH pathway inhibitors but they have not been linked to human HPE, the most common outcome of genetic deficiency in HH signaling (Croen *et al*, 2000; Linn *et al*, 1983; Miller *et al*, 2010; van Gelder *et al*, 2009; van Gelder *et al*, 2010). HPE occurs in ~1:250 conceptions, with 97% succumbing to intrauterine lethality (Leoncini *et al*, 2008; Shiota & Yamada, 2010). Ordinarily, epidemiology would be used to assess whether specific environmental agents are risk factors, but epidemiological studies on possible teratogenic effects of cannabis have neither been designed nor powered to detect a link with HPE. Major reasons for these conclusions are: 1) the 97% in utero lethality rate, leading to low case numbers (Muenke & Beachy, 2001); 2) the window of sensitivity to teratogen-induced HPE in model systems is narrow and equivalent to the third week of human pregnancy (Heyne *et al*., 2015), a time many women do not yet know they are pregnant. Such women may use cannabis at this stage, stop when they know they are pregnant, and self-report as not using the drug (Linn *et al*., 1983); and 3) complex etiologies that involve subthreshold insults, as can occur with HPE, are often difficult to parse with epidemiology. Nevertheless, recent findings that correlate increased cannabis usage with specific structural birth defects (Reece & Hulse, 2019; Reece & Hulse, 2020), combined with our identification of THC as a conditional teratogen, indicates further research into this question is important and of significant public health relevance.

## Materials and methods

### Cannabinoids

THC and CBD were obtained from the NIDA Drug Supply Program. Δ9-THC (50 mg/ml in ethanol solution) was evaporated under nitrogen gas, dissolved in 0.9% NaCl (saline) containing 0.3% Tween 80 to a concentration of 0.75 mg/ml (DiNieri & Hurd, 2012) for administration to mice. THC and CBD were diluted to 0.75 mg/ml in ethanol solution for treatment of cells in vitro.

### Mice

All animal procedures were conducted in accordance with institutional guidelines for the care and use of laboratory animals as approved by the Institutional Animal Care and Use Committee (IACUC) of the Icahn School of Medicine at Mount Sinai. Two to three-month-old *Cdon*^+/-^ mice on a 129S6/SvEvTac (129S6) background (Cole & Krauss, 2003; Hong & Krauss, 2012) were mated for one hour and plugged females were collected. The time of the plug was designated as embryonic day (E) 0.0. THC was injected intraperitoneally at selected times and various doses with 0.3% Tween-80 in saline as a vehicle. Offspring were examined by whole mount at E14.0 for external signs of HPE. Injected females were also assessed for plasma concentration of THC and metabolites over 60 hours using a BIOO Scientific MaxSignal THC ELISA kit, according to the manufacturer’s instructions.

### Whole mount in situ hybridization

Whole mount RNA in situ hybridization was performed according to standard protocols (Hong and Krauss, 2012). Briefly, E10 embryos were dissected out and fixed in 4% paraformaldehyde in phosphate-buffered saline (PBS), dehydrated through a graded methanol series, and stored at −20°C. Rehydrated embryos were treated with 10 μg/ml proteinase K (Qiagen) in PBS, 0.1% Tween-20 (PBST) according to stage. Embryos were rinsed with PBST, post-fixed and hybridized with digoxygenin (DIG)-labeled probe in hybridization mix (50% formamide, 1.3x SSC pH 5.0, 5 mM EDTA, 50 μg/ml yeast tRNA, 100 μg/ml heparin, 0.2% Tween-20, and 0.5% CHAPS) overnight at 65°C. They were washed, blocked with 2% Roche blocking reagent, 20% heat-inactivated lamb serum in Tris-buffered saline with 0.1% Tween-20 (TBST) and incubated with alkaline phosphate-conjugated anti-DIG antibody (1:2000, Roche) in blocking buffer overnight at 4°C. After washing with TBST and NTMT (100 mM NaCl, 100 mM Tris pH9.5, 50 mM MgCl2, and 0.1% Tween-20), embryos were stained with BM Purple AP substrate (Roche) in the dark. Stained embryos were cleared in 80% glycerol and photographed with a Jenoptik ProgRes C3 camera attached to Nikon SMZ 1500 stereomicroscope. Captured images were assembled by Helicon Focus software (Helicon Soft). Whole mount in situ hybridization signal intensity was quantified. Briefly, for all embryos, the channels of the original RGB (red, green and blue) images were separated to analyze the specific emission channel. The mean intensity of in situ hybridization signals within the dashed lines of individual embryos (Figure 1C) were calculated with Fiji Image J software.

### Histology and immunohistochemistry

Embryos were fixed in 4% paraformaldehyde and embedded in paraffin for preparation of 7-μm-thick sections for H&E staining. Sections were then dehydrated through graded ethanol and xylene and mounted with Permount (Fisher Scientific). For VNT staining, embryos were infiltrated with sucrose and then embedded in O.C.T frozen medium. Collect 12-μm-thick cryosections of forelimb region and transfer to Superfrost plus slides. After postfixation in 4% paraformaldehyde, sections were washed for 3 times PBST (0.2% Triton X-100). Primary antibodies and secondary antibodies were used blocking buffer (1X PBST plus 3% BSA) to dilute. Sections were incubated in primary antibodies overnight at 4°C. After washing with PBST, incubated in secondary antibodies for 2 hr at room temperature. The primary antibodies were used: OLIG2 (Millipore Sigma, AB9610), CB1R (Cayman Chemical, 10006590), FOXA2 (DSHB, 4C7), NKX2.2 (DSHB, 74.5A5), ARL13B (NeuroMab, 75-287). Anti-rabbit, Alexa 568 or Alexa 488; anti-mouse IgG1, Alexa 647; anti-mouse IgG2b, Alexa 488; anti-mouse IgG2a, Alexa 594 (Invitrogen, 1:500) were used for secondary antibodies. VECTASHIELD antifade mounting medium (Vector Lab, H-1000) was used to prevent signals from fading. Cryosections were imaged by using Leica SP5 DMI confocal microscope with a 20X objective. For VNT staining, five embryos of each group were analyzed. Between 3 and 5 individual sections per mouse were quantified for DAPI, FOXA2, NKX2.2, and OLIG2 positive cells. Data were processed by Fiji Image J software.

### Plasmids

A mouse *Cnr1* cDNA fragment (Addgene, #13391) was cloned into all-in-one tetracycline inducible plasmid (pAS4.1w.Ppuro-aON, Academia Sinica, Taiwan). Recombinant lentivirus was produced by transfecting HEK293T cells with Lipofectamine 2000 (Invitrogen, 11668019) or TransIT-2020 transfection reagent (Mirus, MIR5404) and a third generation lentiviral package system (Dull *et al*, 1998). Recombinant lentivirus was used to infect NIH3T3 cells and stably expressing cells selected with 5 μg/ml puromycin (Thermo Fisher, A1113803). Conditional expression of CB1R was induced by the addition of Doxycycline (Millipore Sigma, D9891).

### Cell culture and qRT-PCR

NIH3T3 cells, HEK293T, mouse embryonic fibroblasts (MEFs) and Neuro-2a cells were grown in DMEM supplemented with 10% fetal bovine serum, penicillin, and streptomycin. For qRT-PCR, 80% confluent NIH3T3 cells or MEFs were cultured overnight in DMEM plus 2% FBS, after which they were incubated for another 24 hr in fresh DMEM plus 2% FBS supplemented with 5 ng/ml SHH (recombinant human amino terminal fragment of SHH, StemRD) in the presence or absence of desired antagonist. After incubation, total RNA was extracted using RNeasy mini kit (Qiagen). For mouse E14 brain, total RNA was extracted by Trizol/RNeasy mini kit. Reverse transcription and cDNA production were performed with Superscript III first strand synthesis system (Invitrogen). qPCR was performed using iQ SyBR green supermix (BioRad) on an iCycler iQ5 (BioRad). Gene expression levels were normalized to *Gapdh*. The primer sequences are shown as follows:

*Gapdh*, AACGACCCCTTCATTGAC (Forward primer) and TCCACGACATACTCAGCAC (Reverse primer) (Dong *et al*, 2008);
*Gli1*, CCAAGCCAACTTTATGTCAGGG (Forward primer) and AGCCCGCTTCTTTGTTAATTTGA (Reverse primer) (Harvard primer, 6754002a1);
*Cnr1*, CTGGTTCTGATCCTGGTGGT (Forward primer) and TGTCTCAGGTCCTTGCTCCT (Reverse primer) (Ludányi *et al*, 2008).

### Immunofluorescence and ciliary SMO localization measurement

Cells were plated on rat tail collagen (Sigma-Aldrich, 11179179001) coated glass slides. After incubation, cells were fixed in 4% paraformaldehyde at room temperature for 10 min and then in 100% methanol at −20°C for 4 min. Fixed cells were permeabilized in 0.1% Triton X-100 and blocked in 1X PBS plus 3% BSA for 30 min. Cells were incubated in primary antibodies overnight at 4°C and then secondary antibodies for 2 hr at room temperature. The primary antibodies used were SMO (Santa Cruz, sc-166685), ARL13B (ProteinTech, 17711-1-AP), γ-tubulin (Abcam, ab11316), and acetylated tubulin (Sigma-Aldrich, T7451). Anti-rabbit, Alexa 488; anti-mouse IgG1, Alexa 647; anti-mouse IgG2a, Alexa 594 (ThermoFisher, 1:500) were used for secondary antibodies. Images were captured on a Leica SP5 DMI confocal microscope with a 63X oil objective. A shifted overlay of SMO and ARL13B signals was used for better visualization. For quantification of ciliary SMO, first a mask was determined by using the ARL13B image, and then the mask was applied to the corresponding SMO image where the fluorescence intensity was measured. An identical area nearby but outside the cilia was also measured to determine background fluorescence. Background subtraction was applied on each primary cilium. The percentage of ciliary SMO was determined by fluorescence intensity. Compared with the mean value of positive controls, when treatments decreased SMO signal by more than 50%, cilia were considered negative. The lengths of primary cilia were measured by the ARL13B signal. Between 100 and 140 cilia were analyzed per condition. Data were quantified by Fiji Image J software.

### Immunoprecipitation

To detect the interaction of CB1R and SMO, mouse brain lysates wer harvested in lysis buffer containing 50 mM Tris-HCl (pH 8.0), 150 mM NaCl, 2 mM EDTA, 10% glycerol, 0.5% Nonidet P40, 5 mM NaF, 1mM Na_3_VO_4_, and a protease inhibitor cocktail (Sigma–Aldrich). Whole brain lysate was precleared with Protein G agarose (Roche, 11243233001) and the supernatant was immunoprecipitated with CB1R antibody overnight at 4°C. The lysate was then incubated with Protein G agarose for 3 hr at 4°C. Material pulled down was washed with lysis buffer 3 times and eluted by boiling. Eluates were separated by SDS-PAGE and transferred to PVDF membranes which were immunoblotted with primary antibodies, CB1R and SMO, overnight 4°C. Secondary antibodies used for western blotting were: anti-mouse (Cell Signaling, #7076), HRP-conjugated anti-rabbit (Cell Signaling, #7074); or HRP-conjugated anti-rabbit secondary antibodies, light chain specific (Genetex, GTX221666; Jackson Lab, 211-032-171), which do not detect IgG heavy chain.

### BODIPY-Cyclopamine binding assays

This protocol was as detailed by Chen et al. (Chen *et al*., 2002) with minor modifications. Briefly, SMO tagged at its C terminus with Myc epitope was expressed in 293T cells by transient transfection. Empty vector was transfected as a control. Twenty-four hours later, cells were plated on rat tail collagen-coated glass slides, and 24 hours after that were incubated for 4 hr in OPTI-MEM with 5nM BODIPY-Cyclopamine, in the presence or absence of competitor drug at 37°C. The cells were washed with PBS and fixed in 4% paraformaldehyde for 15 min, followed by permeabilization with PBST (containing 0.1 % Triton X-100). SMO was detected with Myc antibody (Santa Cruz, sc-40) and Alexa 568-conjugated mouse secondary antibody (ThermoFisher, 1:500). The cells were then imaged on a Leica DM5500 B upright microscope, capturing 10 fields of view. Representative fields are shown. Cells not expressing SMO-Myc or not incubated with BODIPY-Cyclopamine were used for background subtraction for both BODIPY and Alexa568 channels. Data is presented as background-subtracted BODIPY intensity and Alexa568 intensity for SMO-Myc–expressing cells.

### Statistical analysis

Experiments were done at least three independent times. Data was analyzed with GraphPad Prism 8. Graphs show mean values and error bars indicating standard deviation (SD). Statistical analysis of the data was performed using Student’s t test or two-tailed Fischer’s exact test as noted. Statistical significance was classified as *P<0.05, **P<0.01 or ***P<0.001. A P value of <0.05 was considered statistically significant.

## Data availability

This study includes no data deposited in external repositories.

## Acknowledgements

The authors dedicate this paper to the memory of Suzanne Eaton. We thank Rune Toftgard for providing *Ptch1*^-/-^ and *Sufu*^-/-^ MEFs, Lakshmi Devi for providing Neuro-2a cells, Hungwen Chen for expression vectors, and Paul Wassarman and Phil Soriano for critical comments on the manuscript. This work was supported by a pilot grant from the Mindich Child Health and Development Institute at Mount Sinai and NIH/NIDCR grant DE024748 to R.S.K and by NIH/NIDA grant DA030359 to Y.L.H.

## Author contributions

HFL performed and analyzed the majority of the experiments and prepared the figures. MH performed and analyzed experiments in Figure 1 and helped prepared the figure. HS assisted with experiments using THC in vivo. YLH provided guidance on experiments and cannabinoids. RSK conceived and supervised the study. HFL and RSK wrote the manuscript. All authors edited the manuscript.

---

**Supplemental Table 1.** Frequency of THC-induced HPE in *Cdon*^-/-^ embryos.

| Treatment | # *Cdon*<sup>+/+</sup> or *Cdon*<sup>+/-</sup> embryos | # *Cdon*<sup>+/+</sup> or *Cdon*<sup>+/-</sup> embryos with HPE | # *Cdon*<sup>+/-</sup> embryos | # *Cdon*<sup>-/-</sup> embryos with HPE (%) | P value, Fisher's exact test (vs.VEH) |
| --- | --- | --- | --- | --- | --- |
| Vehicle | 29 | 0 | 15 | 0 |  |
| THC (5 mg/kg) | 33 | 0 | 12 | 2 (16.7) | >0.05 |
| THC (10 mg/kg) | 32 | 0 | 13 | 4 (30.8) | 0.03 |
| THC (15 mg/kg) | 27 | 0 | 11 | 4 (36.4) | 0.02 |

**Supplemental Table 2.**
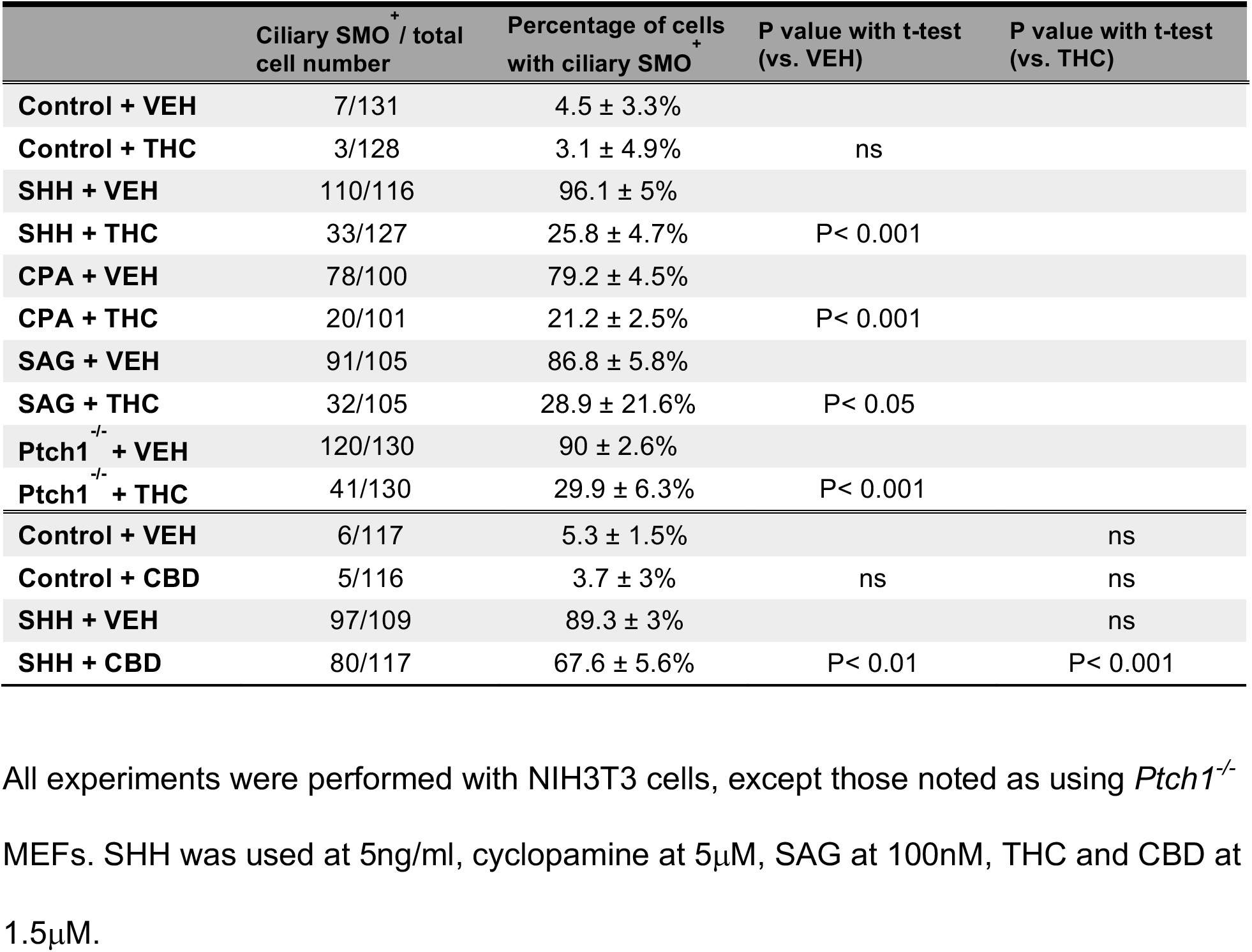
Effects of THC and CBD on SMO localization to primary cilia.

## Supplemental Figure Legends

**Supplemental Fig. 1.**
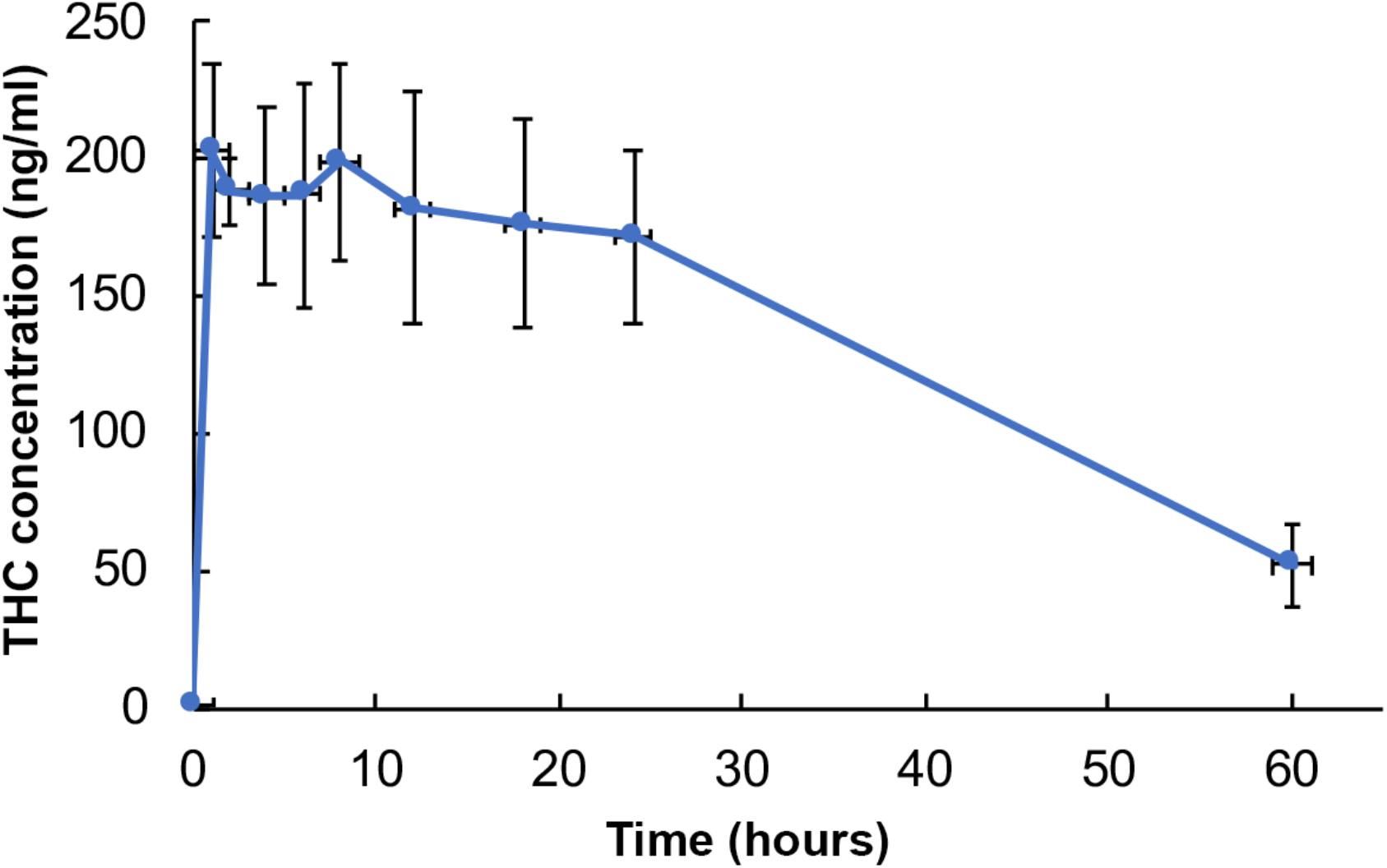
Maternal blood THC levels. The level of THC plus THC metabolites was measured over a 60-hour time course after IP administration of 15 mg/kg THC. Serum was collected and analyzed by ELISA. Values are means ± SD, N=4-10 mice per time point.

**Supplemental Fig. 2.**
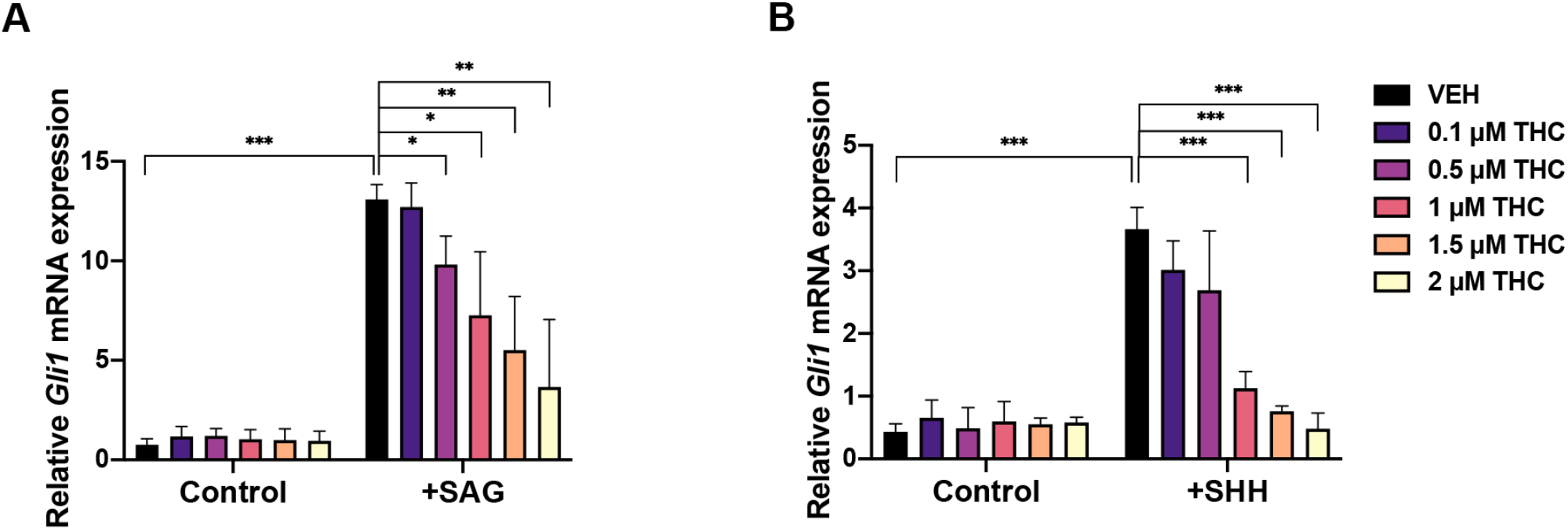
THC suppresses SHH-induced *Gli1* expression in MEFs. A. NIH3T3 cells were treated with 50nM SAG and the indicated concentrations of THC for 24 hr. B. MEFs were treated with 5 ng/ml SHH and the indicated concentrations of THC for 24 hr. Relative *Gli1* mRNA expression was analyzed by qRT-PCR. Values are means ± SD, N=3. *, ***, p<0.05, <0.001 with Student’s t-test.

**Supplemental Fig. 3.**
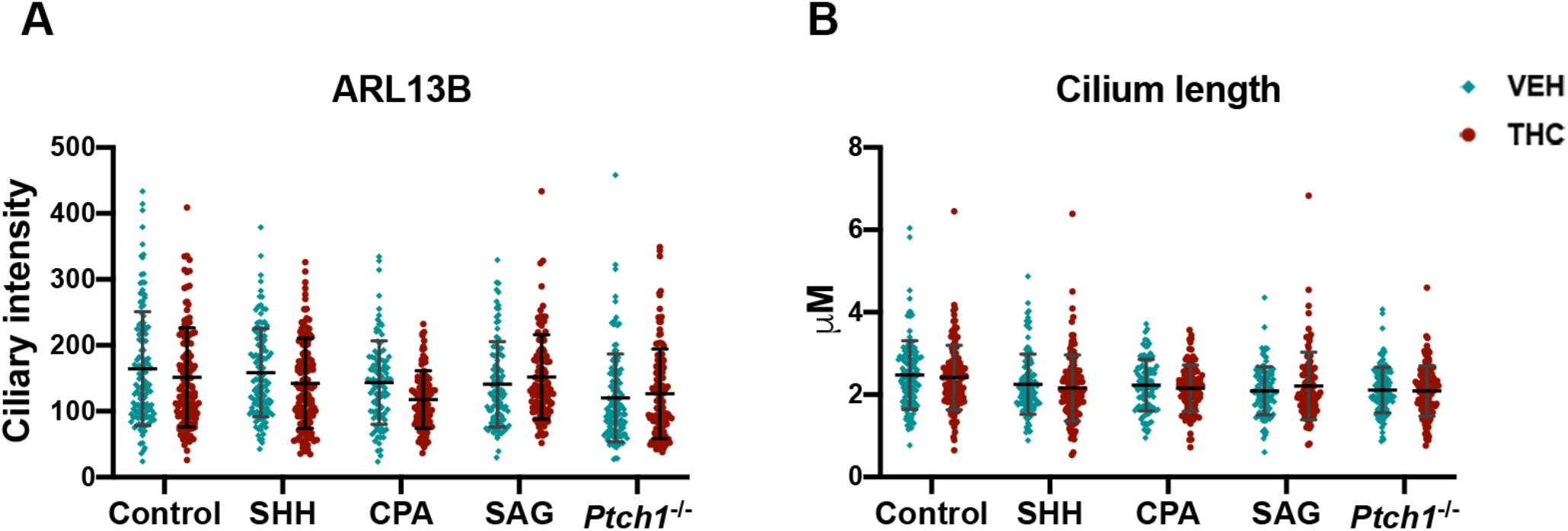
THC does not grossly affect primary cilia. A. Fluorescence intensity for the primary cilia marker, ARL13B. B. Primary cilia length. Each point represents an individual cell collated from at least 3 independent experiments.

**Supplemental Fig. 4.**
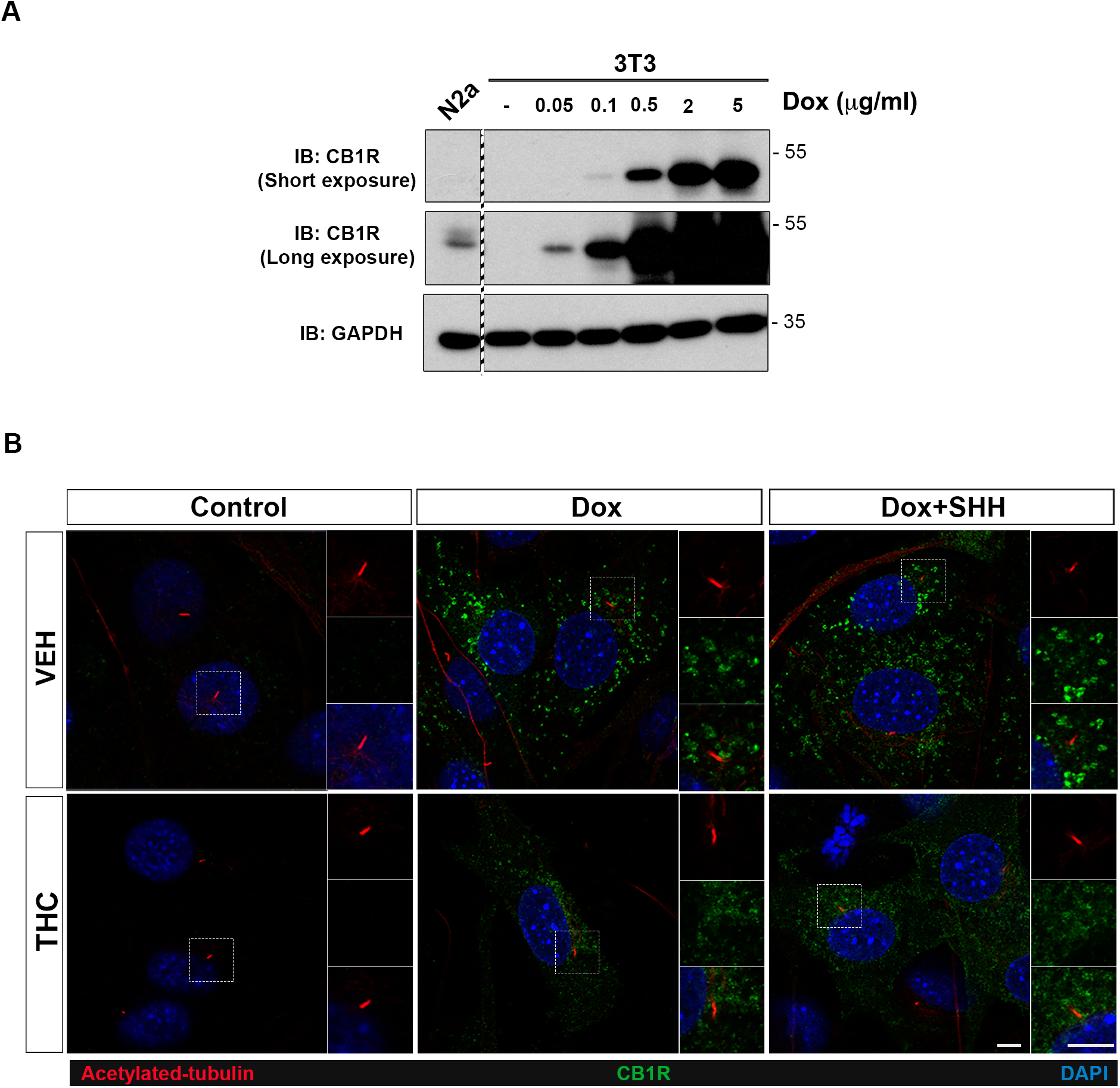
NIH3T3 cells with inducible expression of CB1R. A. NIH3T3 cells were infected with a lentiviral vector encoding doxycycline-inducible expression of CB1R. Cells were treated with the indicated doses of doxycycline (Dox) and harvested for western blot analysis 24 hours later. N2a, Neuro-2a cells, a cell line that expresses CB1R endogenously. The samples are from the same gel and membrane. The dashed region indicates lanes not shown. B. Exogenously expressed CB1R does not localize to primary cilia in NIH3T3 cells, with or without SHH stimulation. Primary cilia are marked by acetylated tubulin. Scale bars, 5μm.

